# A mutation in the promoter of the *pmr*D gene of *Shigella flexneri* abrogates functional PhoPQ-PmrD-PmrAB signaling and polymyxin B resistance

**DOI:** 10.1101/2021.07.26.453917

**Authors:** Raymond Huynh, Joseph B. McPhee

## Abstract

*Shigella* spp. are the causative agent of bacillary dysentery, a major cause of food-borne morbidity and mortality worldwide. These organisms are recently evolved, polyphyletic pathovar of *E. coli*, and since their divergence they have undergone multiple cases of gene gain and gene loss and understanding how gene inactivation events alter bacterial behaviour represents an important objective to be better able to understand how virulence and other phenotypes are affected. Here, we identify a frameshift mutation in the *pmrD* gene of *S. flexneri* that although it would be predicted to make a functional, full-length protein, no such production occurs, likely due to the non-optimal spacing between the translational initiation site and the Shine-Dalgarno sequence. We show that this loss severs the normal connection between the PhoPQ two-component regulatory system and the PmrAB two-component regulatory system, abrogating low Mg^2+^ mediated cationic antimicrobial peptide and polymyxin B resistance, while maintaining normal PmrAB-mediated polymyxin B resistance. In contrast, *S. sonnei* maintains a functional PmrD protein and canonical signaling through this regulatory network. This species specific gene loss suggests that *S. flexneri* and *S. sonnei* have evolved different regulatory responses to changing environmental conditions.

## Introduction

In order to respond to their environment, bacteria have evolved numerous families of regulatory proteins. One particularly successful family of regulators is the two-component regulatory system. These typically consist of a membrane-bound sensor kinase that responds to a particular environmental stimulus and a cytoplasmic response regulator that mediates a specific output. In most two-component systems, the response regulator is a DNA-binding protein that binds to specific DNA motifs in the promoter of a target gene to increase transcription of those genes by increasing recruitment of RNA polymerase (1). Two component regulatory systems can alter gene expression in response to altered temperature, pH, extracellular ionic composition, specific small molecules or even redox status or light levels (2). Two-component systems can be a simple system controlling one or a few target genes or can evolve into complex phosphotransfer relays that modulate complex changes in cellular physiology like sporulation or virulence (3).

In the Enterobacterales, the PhoPQ system is very important for intestinal fitness, for resistance to HDPs, and for virulence (4–6). The PhoPQ system is a master regulator of virulence in *Salmonella enterica* (5, 7, 8), in *Yersinia pestis* (4, 9–13), and in numerous types of *E. coli*, including UPEC, EPEC and EHEC (14–16). Although the specifics of how PhoPQ contributes to virulence can vary from species to species, and even from strain to strain (6, 17), in general, the PhoPQ system is a key component of a larger signaling network in each of these species of bacterial pathogen. In *E. coli, S. enterica* and *K. pneumoniae*, the PhoPQ system transcriptionally regulates a gene encoding a small protein, PmrD, that post-transcriptionally regulates another two-component system, PmrAB (BasRS) (18–20). The PmrAB system responds directly and independently to extracellular iron but when PmrD is expressed, control of PmrAB-regulated target genes is brought under the control of the PhoPQ system (21, 22). PmrA, in turn, directly regulates numerous genes controlling the structure and behaviour of bacterial lipopolysaccharide, including the *arnBCDATEF* operon (23, 24), the *pmrCpmrAB* operon, and several core and lipid A phosphoethanolamine transferases like *cptA, eptA, eptB* and *eptC* (25, 26). The net effect of these changes is the production of a bacterial outer membrane with reduced anionic charge and therefore reduced interactions with cationic antimicrobial host-defense peptides.

PmrD is a small protein that is required for PhoPQ-mediated regulation of polymyxin B and HDP resistance (21, 27). The solution structure of PmrD is principally composed of a six-stranded antiparallel β-sheet and a C-terminal α-helix (28). Using BeF^3-^ labelled PmrA as a mimic for phospho-PmrA, Luo et al showed that as with other active response regulators, the phosphorylation of PmrA results in dimerization of the receiver domain (28). Both X-ray and NMR mediated structural characterization of PmrD show that it forms a dimeric structure via intermolecular salt bridges. The dimeric PmrD binds to phospho-PmrA with 1:1 stoichiometry (28, 29) and prevents the dephosphorylation of the PmrA receiver domain by the cognate sensor kinase PmrB, thereby explaining the mechanism of PmrD-mediated polymyxin B and HDP resistance (30). Thus, PhoPQ transcriptional control of PmrD brings HDP resistance genes under the control of PhoPQ activating signals (i.e. HDPs).

This mechanism of PhoPQ-PmrD-PmrAB control of HDP resistance is conserved in *E. coli, S. enterica* and *K. pneumoniae*, and is important for intestinal pathogenesis in these organisms. However, the role of the PmrD-PmrAB system in *Shigella* spp. has not been well-described. *Shigella* spp. are a recently evolved, polyphyletic pathovar that are rooted within the *Escherichia* lineage. The PhoPQ system of *Shigella* is required for virulence in the Sereny model of infection as well as for resistance to antimicrobial HDPs (5, 31). Unlike other species of Enterobacterales however, high level *Shigella* HDP resistance depends on the activity of a virulence plasmid encoded lipid A acyltransferase, MsbB2 (32–34). MsbB2 is required for complete acyl-oxyacylation of lipid A and this modification is required for maximal induction of TNF-α production by human monocytes and for inflammatory destruction of the rabbit intestinal epithelial barrier, as well as for full virulence in a murine pneumonia challenge model, indicating that MsbB2-controlled lipid A alterations drive inflammatory processes that lead to maximum virulence (32, 34). The gene for this activity is encoded in an operon found on the large virulence plasmid called *shf-rfbU-virK-msbB2* that is transcriptionally regulated by the PhoP regulatory system (33). This operon also contains the gene *shf*, which encodes SfPgdA, an enzyme that modifies *Shigella* peptidoglycan by deacetylating it, and *rfbU*, encoding SfGtr4, an LPS glucosyltransferase, both resulting in resistance to host-derived lysozyme (35, 36) as well as an orthologue of *Salmonella* VirK, a membrane protein associated with resistance to HDPs (37). Thus, PhoP-regulation of the *shf-rfbU-virK-msbB2* operon appears to control susceptibility to two different types of host defense molecules, HDPs and neutrophil derived lysozyme.

Here, we describe a mutation in the *pmrD* locus of *Shigella flexneri*, whereby the PmrD protein is not produced due to non-optimal spacing between the ribosome binding site and the in-frame start codon. We show that this mutation results in loss of low Mg^2+^ induced polymyxin B resistance, a bacterially derived cationic antimicrobial peptide antibiotic. Restoration of *pmrD*_*Ec*_ to *S. flexneri* restores Mg^2+^ regulated polymyxin B resistance. In contrast, *S. sonnei* maintains a functional *pmrD* gene and responds to PhoPQ-activating signals by increasing resistance to polymyxin B in a *pmrD*-dependent manner. This suggests that these two closely related species of bacteria may experience different environmental stressors during their life cycles such that *S. sonnei* retains PhoPQ-regulated HDP resistance while *S. flexneri* either no longer requires this or *pmrD*-mediated regulation of PmrAB-target genes is maladaptive. Further study of this is warranted due to the significant public health burden associated with *Shigella* spp. infection.

## Materials and methods

### Bacterial strains, plasmids and growth conditions

All strains, plasmids and primers used in this study are shown in Table 1. For the *E. coli* unmarked mutants, appropriate strains were obtained from the Coli Genetic Stock Center at Yale University and their kanamycin resistance cassette was removed by FLP-recombinase mediated expression as described by Datsenko and Wanner (38). For PmrD expression plasmids, synthetic sequences were ordered with the sequences shown in Supplemental Figure 1. Briefly, we designed plasmids that would have conservative frameshifts to bring the full length *pmrD* gene into frame with either start site 1, 2 or 3. All constructs were expressed from the native *E. coli pmrD* promoter. These genes were ordered from BioBasic gene synthesis and subcloned to pWSK129. *E. coli* and *Shigella* spp. were routinely cultured in lysogeny broth at 30°C. When used for functional assays, *Shigella* were first streaked to trypticase soy agar containing 0.01% Congo Red (CR) and incubated overnight at 37°C. CR+ colonies were picked and grown to stationary phase in lysogeny broth at 30°C. Subcultures were then grown in N-minimal medium with 0.2% glucose as a carbon source and supplemented with 0.1% casamino acids (39). Where indicated, medium was supplemented to either 10 mM (high Mg^2+^) or 10 μM (low Mg^2+^) or either 100 μM (high Fe^3+^) or 0.1 μM (low Fe^3+^) to selectively stimulate either the PhoPQ or the PmrAB regulatory systems.

**Table 1:**
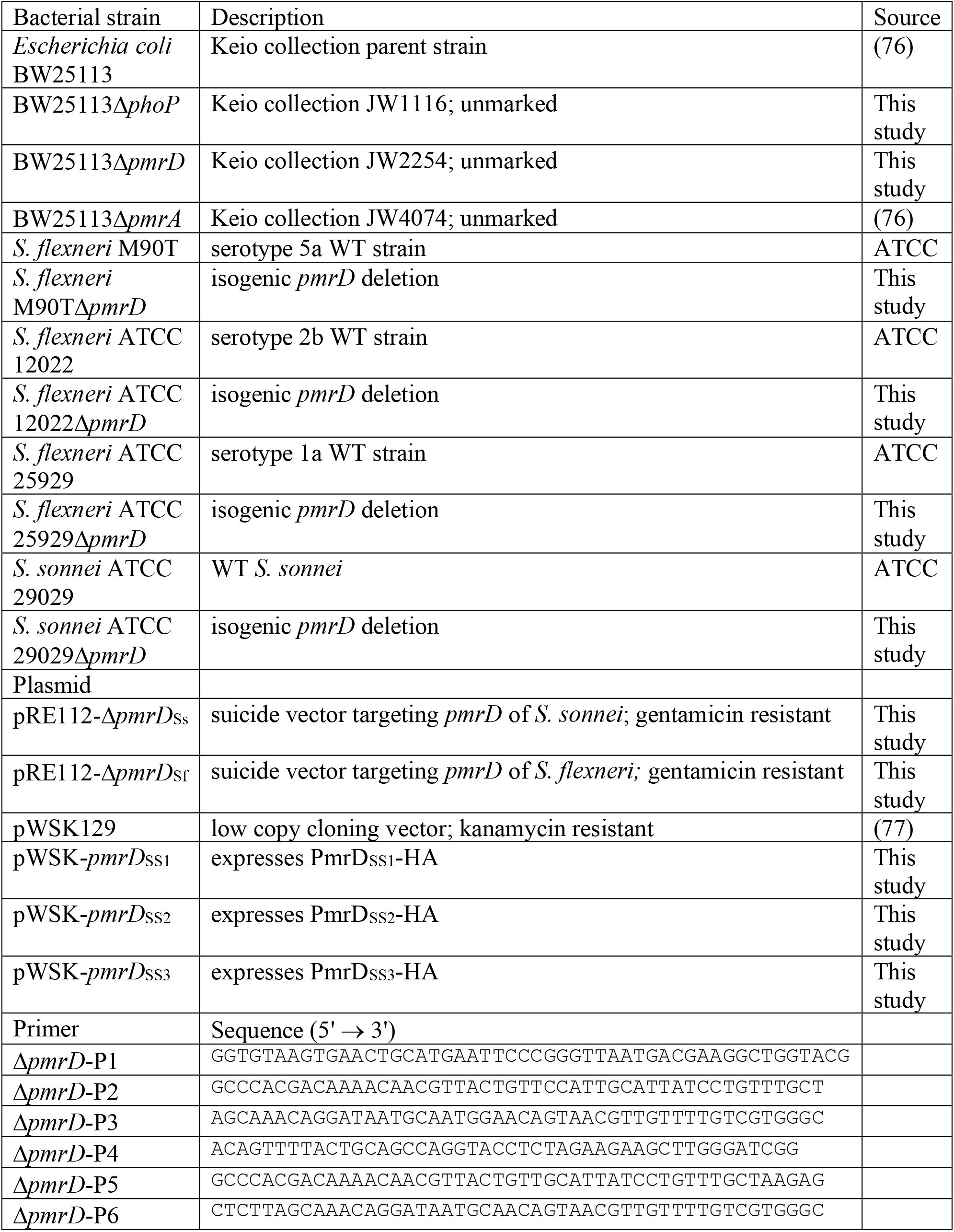
Strains, plasmids and primers used in this study

### Deletion of *pmrD* from *Shigella* spp

The *pmrD* genes of *S. flexneri* and *S. sonnei* were deleted via suicide plasmid mediated allelic exchange. Briefly, a deletion allele consisting of the 500 bp region upstream of *pmrD* and the first two codons along with the final two codons and the downstream 500 bp region was created using strand overlap extension PCR. Different combinations of primers were used to target each strain, depending on the sequence of *pmrD* in that strain. For *S. sonnei, pmrD*-P1, *pmrD*-P2, *pmrD*-P3 and *pmrD*-P4 were used to construct the appropriate targeting cassette while for *S. flexneri, pmrD*-P1, *pmrD*-P5, *pmrD*-P6 and *pmrD*-P4 were used. These fragments were cloned to pCR2.1-TOPO, sequenced, then subcloned to pRE112-Gm, a gentamicin resistant version of pRE112 (40). Biparental mating between S17-1λpir containing pRE112-Δ*pmrD*_Sf_ or pRE112-Δ*pmrD*_Ss_ and *S. flexneri* or *S. sonnei* strains containing the temperature sensitive pKD46 plasmids (38) was carried out. Selection for the recombinant Δ*pmrD* deletion strains was conducted by sequential screening for *Shigella* spp. strains that were both ampicillin and gentamicin resistant. Counterselection on sucrose was then carried out and colony PCR on candidate colonies followed by Sanger sequencing was performed to verify that strains were correct.

### Polymyxin B killing assay

Overnight cultures were pelleted and resuspended in N-minimal media then subcultured 1:25 in N-minimal media containing high/low Mg^2+^ or high/low Fe^3+^ at 37°C with shaking until the cultures reached mid-log growth phase (OD_600_ of 0.4 to 0.7). One mL of culture was pelleted via centrifugation for two minutes at 17,000 × g. The supernatant was discarded, and the cell pellet was resuspended in one mL of sterile 10 mM HEPES buffer pH 7.2 and then normalized to 10^8^ CFU/mL. The normalized cultures were then subjected to polymyxin B treatment. Each 5 µL culture was treated with an equal volume of 1 × PBS or 5 µg/mL, 2.5 µg/mL, or 1.0 µg/mL of polymyxin B (final concentrations: 2.5 µg/mL, 1.25 µg/mL, and 0.5 µg/mL respectively). At t=0, the treated sample was diluted 1:250 in 1x PBS in order to stop killing. Next, the diluted sample was further diluted 1:10 in 1x PBS for a final dilution of 10^−1^. Both the undiluted and 10^−1^ samples were plated onto LB agar plates in four replicates of 10 µL. This step was repeated at 10 minute and 60 minute time points. The plates were then incubated at 20°C overnight and incubated at 37°C for 5 hours prior to counting. Colonies were counted and percentage survival was calculated and analyzed.

## Results

### *S. flexneri* exhibits altered environmental control of polymyxin B resistance compared to *E. coli* and *S. sonnei*

We grew three different strains of *S. flexneri*, one strain of *S. sonnei*, and a K-12 strain of *E. coli* under conditions that would either induce the PhoPQ system (low Mg^2+^, low Fe^3+^), induce the PmrAB system independently of PhoPQ (high Mg^2+^, high Fe^3+^) or that would repress both of these systems (high Mg^2+^, low Fe^3+^). As shown in Figure 1A, we observed that all three strains of *S. flexneri* behaved differently from *E. coli* and *S. sonnei*, with growth in low Mg^2+^ showing no increased polymyxin B resistance compared to growth in high Mg^2+^. We reasoned that this could be due to non-functional PmrD or due to other, unknown alterations in the function of the PmrAB regulon. To test this, we grew *E. coli* and *Shigella* spp. in medium containing high Fe^3+^ and high Mg^2+^, conditions that activate the PmrAB system independently of PhoPQ and PmrD (41). Growth in high Mg^2+^/high Fe^3+^ lead to significantly increased polymyxin B resistance compared to those grown in high Mg^2+^/low Fe^3+^ (Figure 1B). In contrast, we observed low Mg^2+^ and high Fe^3+^ mediated polymyxin B resistance in *E. coli* and *S. sonnei*, (Figure 1B). These observations are consistent with a model whereby *S. flexneri* has severed the connection between PhoPQ and PmrAB.

**Figure 1:**
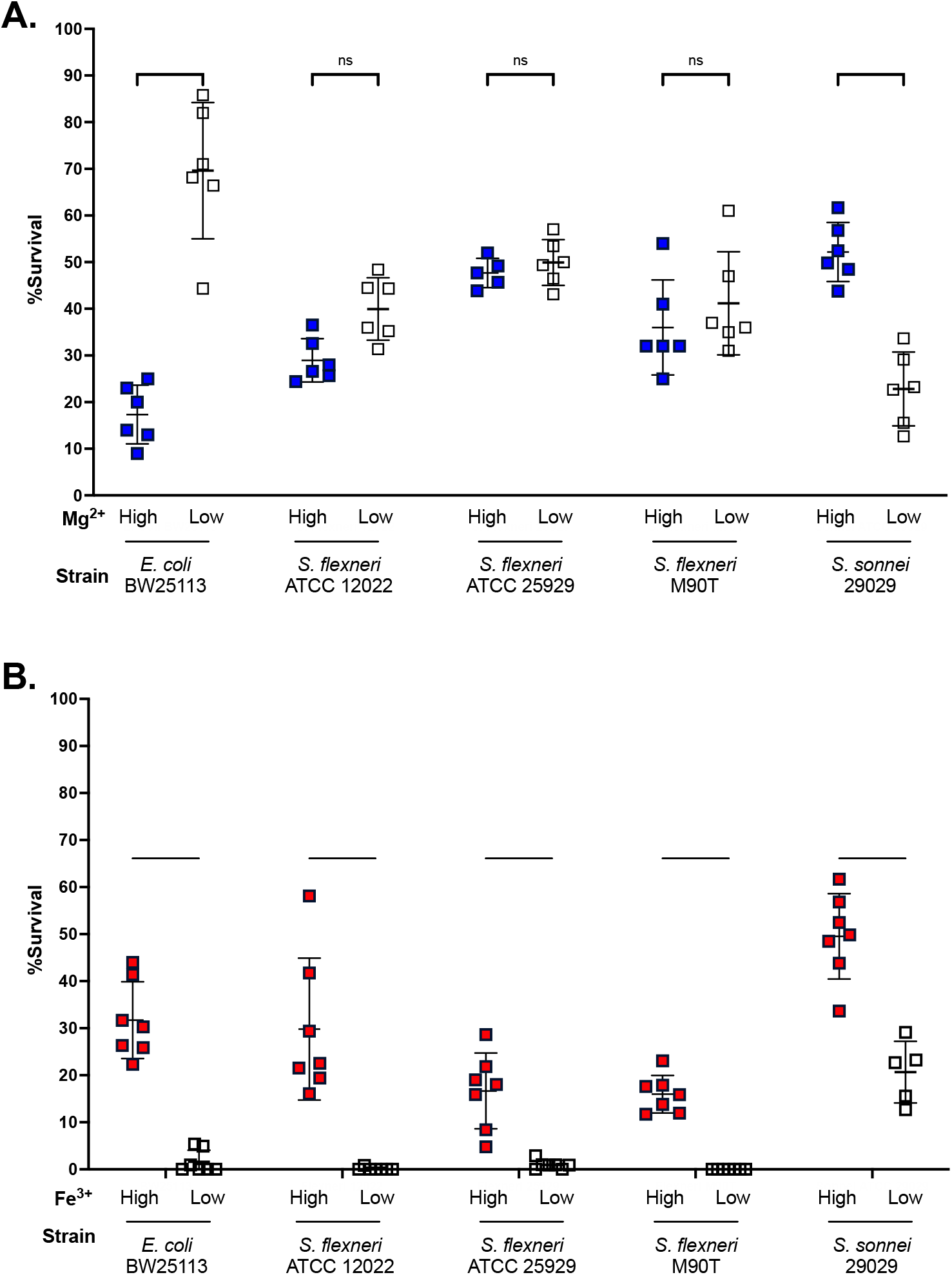
Mg^2+^ dependent regulation of polymyxin B resistance in *Shigella flexneri, S. sonnei* and *E. coli* A) *E. coli* BW25113, *S. flexneri* or *S. sonnei* were grown to mid-log phase in M9-glucose media containing either 1 mM or 10 μM Mg^2+^ and exposed to 1.25 μg/ml polymyxin B for 10 minutes. Surviving bacteria were counted and normalized to bacteria that were not exposed to polymyxin B. B). *E. coli* BW25113, *S. flexneri* or *S. sonnei* were grown to mid-log phase in M9-glucose media containing either 100 μM Fe^3+^ or no added Fe^3+^ and exposed to 1.25 μg/ml polymyxin B for 10 minutes. Surviving bacteria were counted and normalized to bacteria that were not exposed to polymyxin B and the percentage survival was calculated. Each experiment was performed on 5-7 independent biological replicates. ns - p>0.05; ^*^ p<0.05; ^**^ p<0.01; ^***^ p<0.001; ^****^ p<0.0001 by 2-way ANOVA with Šídák’s multiple comparisons test.

### The *pmrD* gene of *E. coli, S. flexneri* and *S. sonnei* contains a start site array sampling three different potential open reading frames

Protein translation in bacteria begins when three initiation factors, a formyl-methionine tRNA and the 30S ribosome itself assemble into a translation initiation complex. The Shine-Dalgarno (SD) sequence of the mRNA interacts directly with the anti-SD sequence of the 30S ribosomal subunit. This interaction brings the initiation codon into the proper site to allow the fMet-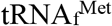 complex to enter the P-site of the nascent ribosome and initiate translation. In bacteria, ∼80% of start codons are ATG, with GTG and TTG making up the remainder (∼12% and 8% respectively)(42). The spacing between the SD and the initiation codon is a crucial factor in the production of an optimal amount of protein (43). Examination of the *pmrD* gene of *E. coli* and *S. flexneri* showed that although they are highly conserved (99.2% identity) the *S. flexneri pmrD* contains a 1 bp insertion in the ORF, which would be expected to bring the coding sequence out of frame with respect to the *E. coli pmrD* start codon. In both *E. coli* and *S. flexneri*, the 5’ end of the gene contains an array of ATG sites, each of which would be predicted to sample a different reading frame (Figure 2). In *E. coli*, the PmrD protein is translated from the second putative start site while in *S. flexneri*, a full-length PmrD protein will only be produced if translation is initiated at the first start site. This suggested that this alteration in spacing between the SD sequence and the translation initiation codon in the mRNA might affect the ability to produce functional PmrD capable of blocking PmrB-mediated dephosphorylation of PmrA.

**Figure 2.**
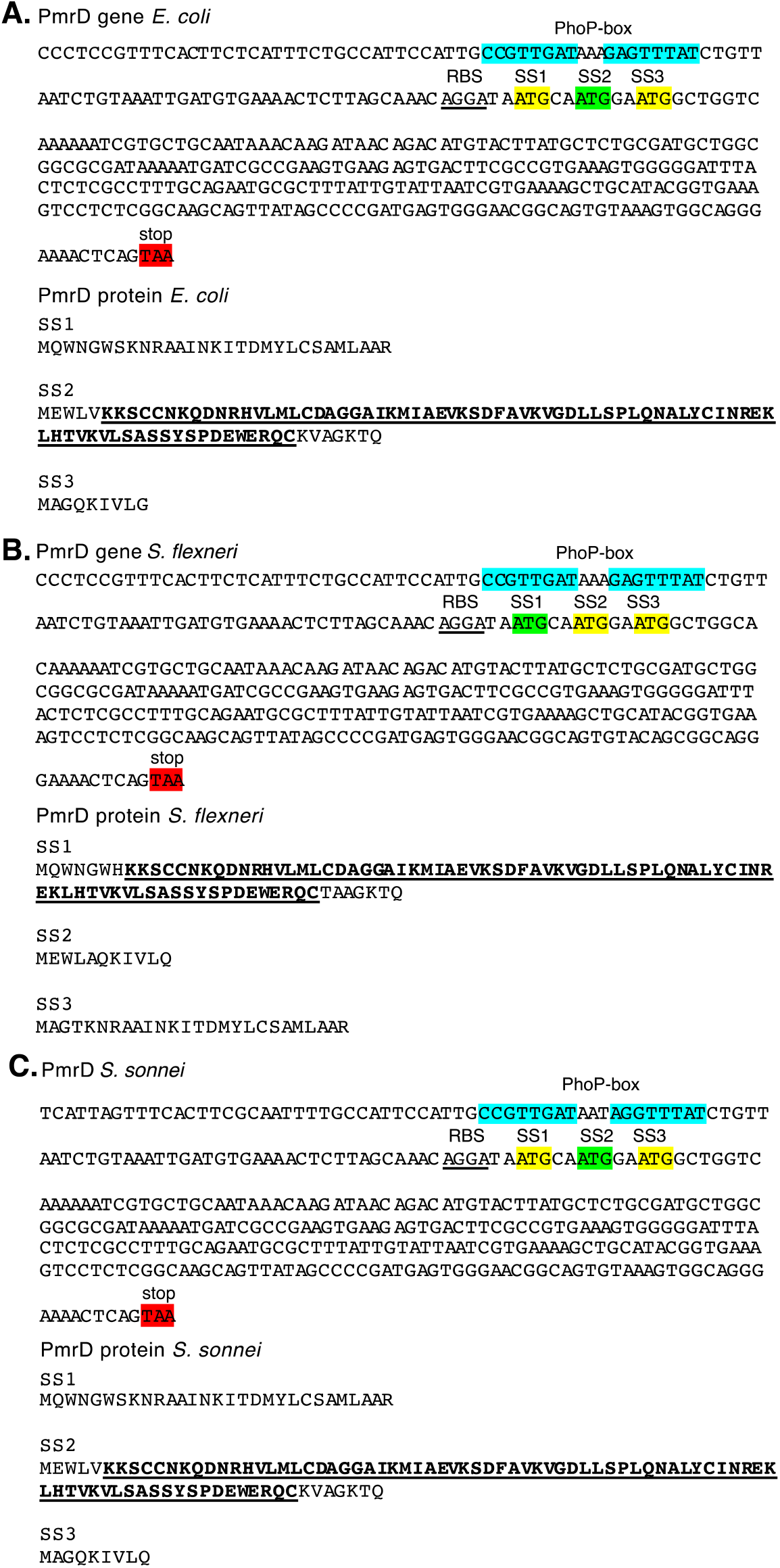
Genes of *pmrD* from (A) *Escherichia coli* (B) *Shigella flexneri* and (C) *Shigella sonnei*. Cyan highlighting shows the PhoP box while green and yellow show the start site arrays in each of these organisms. Green indicates the start site that would produce a full-length and presumably functional PmrD protein. Also shown are the translated proteins that would result if translation were initiated at each of the translational start sites.

### Production of PmrD protein is only observed when translation is initiated at SS2

In order to measure the production of PmrD protein, an HA-epitope tag was fused to three different BW25113 *pmrD* genes representing the SS1, SS2 and SS3 translational start sites followed by western blotting using an anti-HA antibody (Supplemental Figure 1). Each of these alleles was expressed from the low copy pWSK129 plasmid. We transformed the Keio collection BW25113Δ*pmrD* strain with pWSK*pmrD*_SS1_, pWSK*pmrD*_SS2_, or pWSK*pmrD*_SS3_ and protein production was quantified relative to the control protein, DnaK (44) following growth in N-minimal medium containing high (2 mM) or low (20 μM) Mg^2+^. As seen in **Figure 3**, expression was greatest in BW25113Δ*pmrD* + pWSK*pmrD*_SS2_ grown under low Mg^2+^ conditions, while no expression was observed from pWSK*pmrD*_SS1_ or pWSK*pmrD*_SS3_. Consistent with previous reports, PhoP was required for maximal PmrD_SS2_ production, as expression in BW25113 *phoP* strains produced no PmrD-HA, indicating that production of plasmid-encoded PmrD was regulated by the PhoPQ system, as reported previously (22, 27). This is consistent with a model whereby the spacing between the SD and the translational start site is an important determinant of whether or not functional PmrD protein is produced in a given strain of *Shigella* spp.

**Figure 3:**
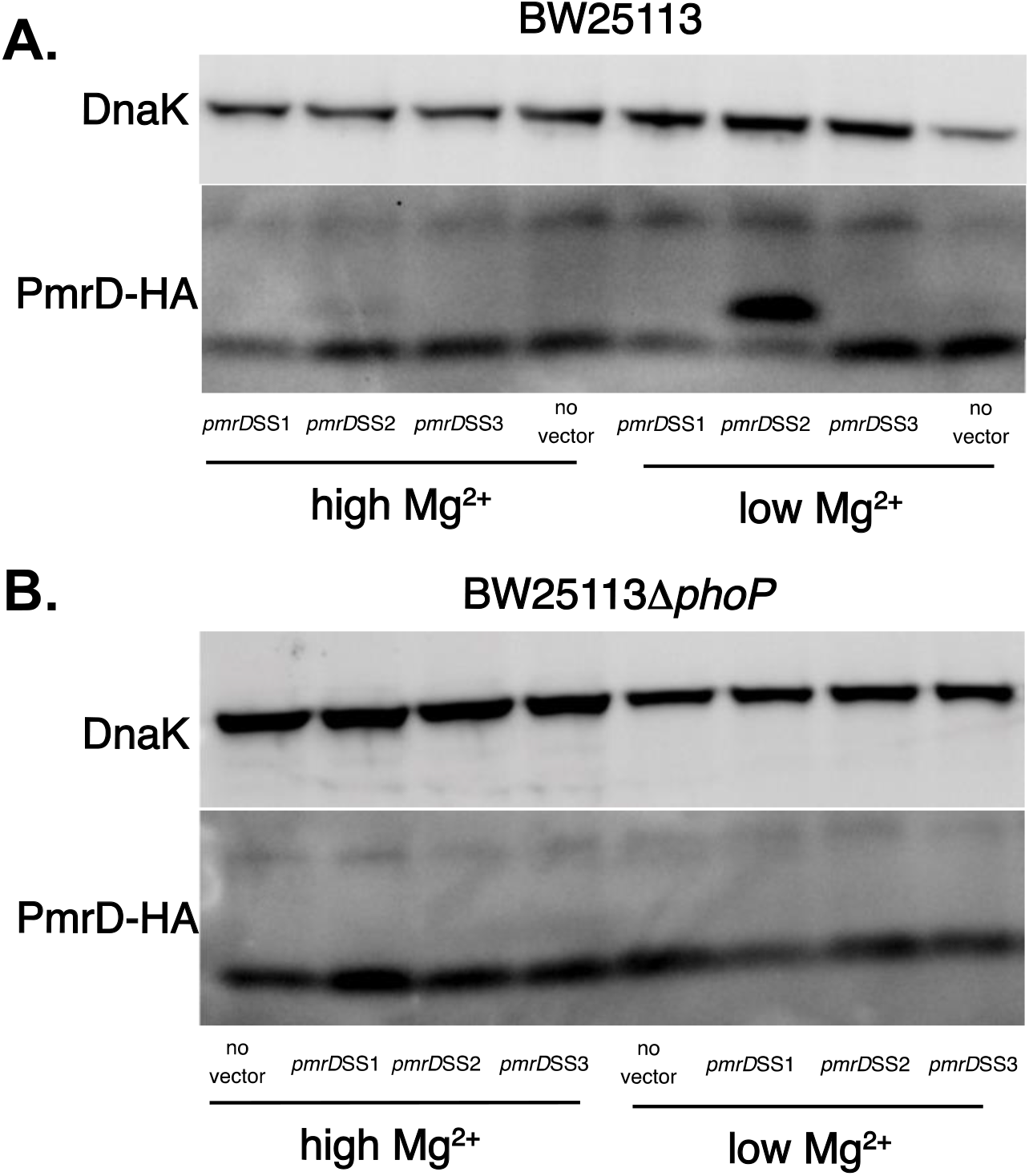
Mg^2+^ regulated PmrD expression of PmrD_SS1_, PmrD_SS2_ and PmrD_SS3_ A) Western blot of PmrD_SS1_, PmrD_SS2_ or PmrD_SS3_ expressed in *E. coli* BW25113 in M9 glucose containing high (2 mM) or low (20 μM) Mg^2+^. DnaK was blotted as a loading control. B) Western blot of PmrD_SS1_, PmrD_SS2_ or PmrD_SS3_ expressed in *E. coli* BW25113Δ*phoP* in M9 glucose containing high (2 mM) or low (20 μM) Mg^2+^. DnaK was blotted as a loading control.

### PmrD_SS2_, but not PmrD_SS1_ or PmrD_SS3_, is required for low Mg^2+^ mediated polymyxin B resistance

To verify that the different spacing arrangements we observed could alter the function of the encoded PmrD protein, we expressed plasmid encoded HA-tagged PmrD proteins in which the *E. coli pmrD* CDS was expressed as a full-length product from either SS1, SS2 or SS3 (Supplemental figure 1). Killing assays were conducted on a BW25113Δ*pmrD* strain containing either no vector or each of the pWSK*pmrD* alleles in addition to BW25113*phoP*, BW25113Δ*pmrD* or BW25113Δ*pmrA* mutants that do not contain a *pmrD*-expressing plasmid. As shown in **Figure 4A**, Mg^2+^-dependent polymyxin B resistance depended on the presence of intact *phoP, pmrD* and *pmrA*. Furthermore, pWSK*pmrD*_SS2_ was only able to complement the resistance phenotype of a BW25113Δ*pmrD* mutant and not that of BW25113Δ*phoP* or BW25113Δ*pmrA* (**Figure 4B**).

**Figure 4.**
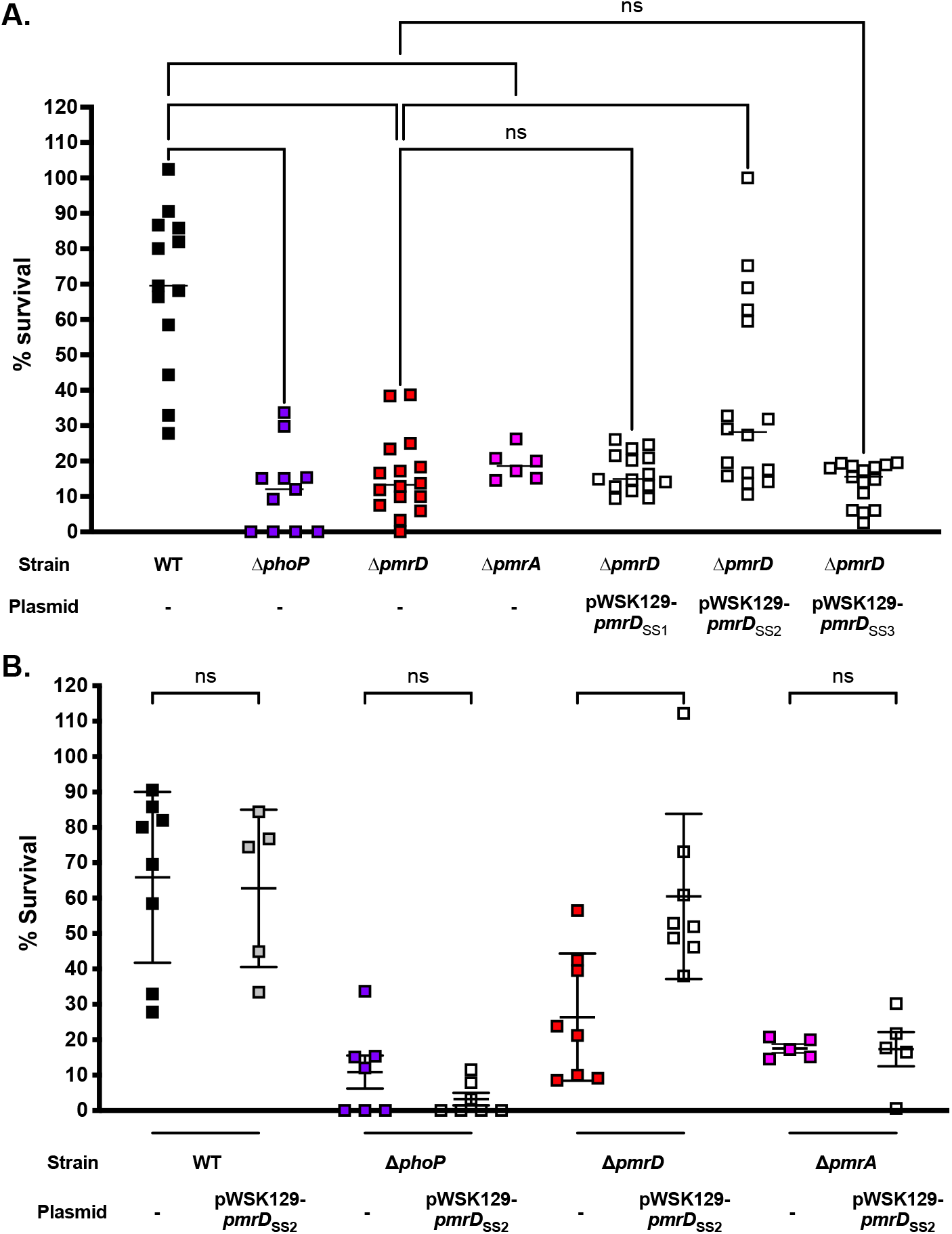
Complementation of a *phoP, pmrD* and/or *pmrA* mutants of BW25113 with PmrD_SS1_, PmrD_SS2_ or PmrD_SS3_. A) BW25113, BW25113 Δ*phoP*, BW25113 Δ*pmrD*, BW25113 Δ*pmrA* and BW25113Δ*pmrD* containing pWSK*pmrD*_SS1_, pWSK*pmrD*_SS2_ or pWSK*pmrD*_SS3_ were grown to mid-logarithmic phase in M9 glucose medium containing 10 μM Mg^2+^, then washed and exposed to 1.25 μg/ml polymyxin B for 10 minutes. Surviving bacteria were normalized to bacteria that were not exposed to polymyxin B and these values were calculated as a percentage survival. B) BW25113, BW25113Δ*phoP*, BW25113Δ*pmrD*, BW25113Δ*pmrA* containing no vector or pWSK*pmrD*_SS2_ were grown to mid-logarithmic phase in M9 glucose medium containing 10 μM Mg^2+^, then washed and exposed to 1.25 μg/ml polymyxin B for 10 minutes. Surviving bacteria were normalized to bacteria that were not exposed to polymyxin B and these values were calculated as a percent survival. Each experiment was performed on 6-16 independent biological replicates. ns - p>0.05; ^*^ p<0.05; ^**^ p<0.01; ^***^ p<0.001; ^****^ p<0.0001 by 2-way ANOVA with Šídák’s multiple comparisons test.

### Plasmid-encoded *pmrD*_SS2_ restores Mg^2+^ induced polymyxin B resistance to *S. flexneri*

To further establish that the *E. coli* PmrD allele is necessary and sufficient to restore normal PhoPQ-PmrD-PmrAB functioning to *S. flexneri*, we constructed *pmrD* deletion mutants of three strains of *S. flexneri* (M90T, ATCC 25929 and ATCC 12022) as well as *S. sonnei* strain ATCC 25931. We then examined the behaviour of both strains when grown under PhoPQ- or PmrAB-inducing conditions. As seen in Figure 5A and 5B, *pmrD* mutants *S. flexneri* do not exhibit any change in Mg^2+^-induced polymyxin B resistance. In contrast, deletion of *pmrD* from *S. sonnei* results in the loss of low Mg^2+^ induced polymyxin B resistance, consistent with *S. sonnei* maintaining a functional PhoPQ-PmrD-PmrAB signaling system (Figure 5B). Complementation of *S. flexneri* and *S. sonnei* with plasmid-encoded *pmrD*_SS2_ results in restoration of low Mg^2+^ induced polymyxin B resistance. All *Shigella* spp WT, *pmrD* and complemented strains tested exhibit normal, Fe^3+^ induced polymyxin B resistance (Figure 5C and 5D).

**Figure 5.**
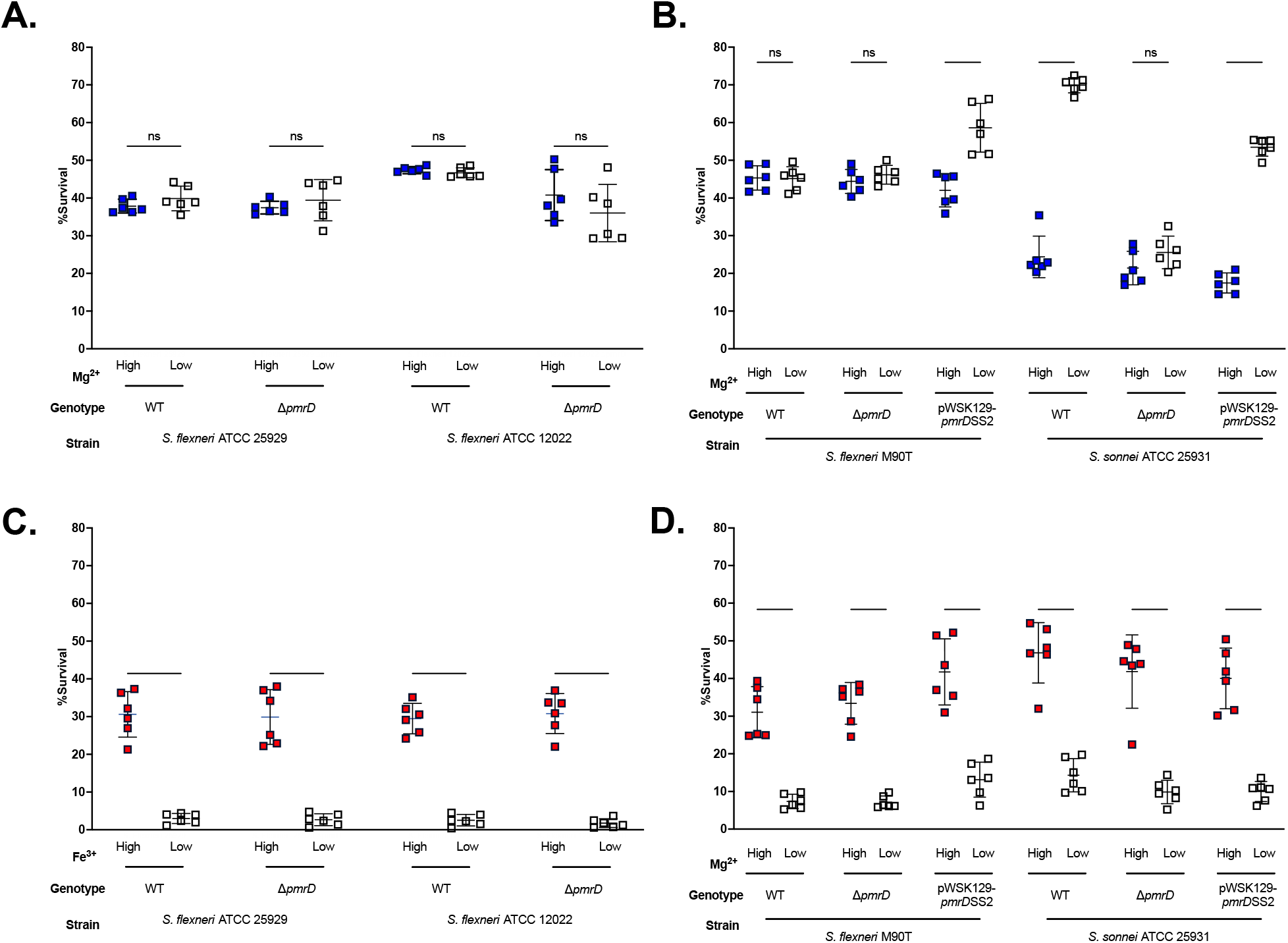
Effect of *pmrD* deletion on Mg^2+^ dependent and Fe^3+^ dependent regulation of polymyxin B resistance. A) *S. flexneri* or isogenic Δ*pmrD* mutants of those strains were grown to mid-logarithmic phase in M9-glucose medium containing 1 mM or 10 μM Mg^2+^, then washed and exposed to 1.25 μg/ml polymyxin B for 10 minutes. Surviving bacteria were normalized to bacteria that were not exposed to polymyxin B and these values were calculated as a percentage survival. B) *S. flexneri* M90T and *S. sonnei* ATCC 25931, isogenic Δ*pmrD* mutants of those strains and the isogenic Δ*pmrD* mutants complemented with pWSK129-*pmrD*_SS2_ were grown to mid-logarithmic phase in M9-glucose medium containing 1 mM or 10 μM Mg^2+^, then washed and exposed to 1.25 g/ml polymyxin B for 10 minutes. Surviving bacteria were normalized to bacteria that were not exposed to polymyxin B and these values were calculated as a percentage survival. C) *S. flexneri* or isogenic Δ*pmrD* mutants of those strains were grown to mid-logarithmic phase in M9-glucose medium containing either no added Fe^3+^ or 100 μM Fe^3+^, then washed and exposed to 1.25 μg/ml polymyxin B for 10 minutes. Surviving bacteria were normalized to bacteria that were not exposed to polymyxin B and these values were calculated as a percentage survival. D) *S. flexneri* M90T and *S. sonnei* ATCC 25931, isogenic Δ*pmrD* mutants of those strains and the isogenic Δ*pmrD* mutants complemented with pWSK129-*pmrD*_SS2_were grown to mid-logarithmic phase in M9-glucose medium containing either no added Fe^3+^ or 100 M Fe^3+^, then washed and exposed to 1.25 g/ml polymyxin B for 10 minutes. Surviving bacteria were normalized to bacteria that were not exposed to polymyxin B and these values were calculated as a percentage survival. Each experiment was performed on 5-7 independent biological replicates. ns - p>0.05; ^*^ p<0.05; ^**^ p<0.01; ^***^ p<0.001; ^****^ p<0.0001 by 2-way ANOVA with Šídák’s multiple comparisons test.

**Figure 6:**
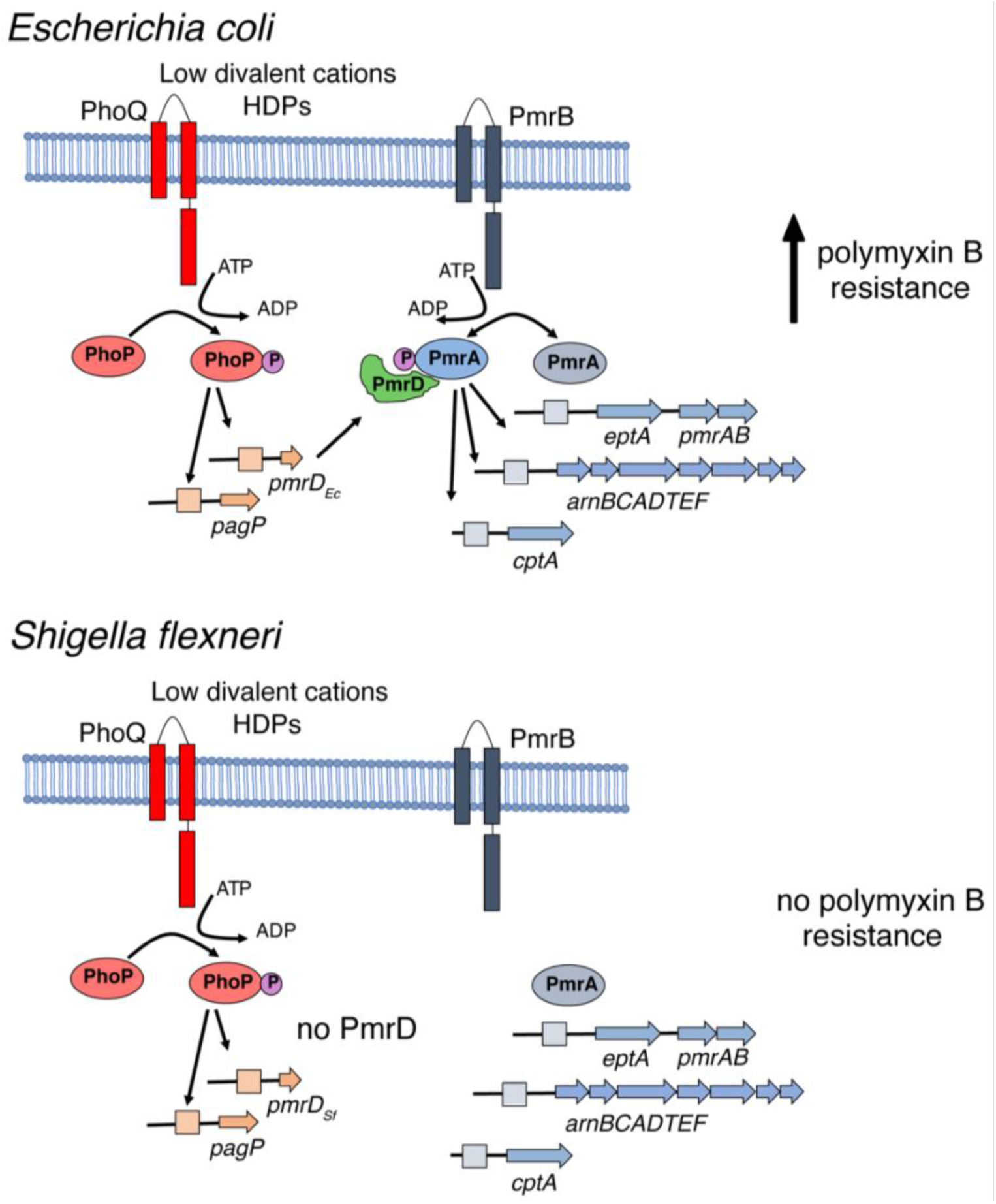
Model showing how the lack of functional PmrD in *Shigella flexneri* results in the loss of PhoPQ-dependent polymyxin B resistance.

## Discussion

*Shigella* spp. are a polyphyletic genus of bacteria, that have evolved on numerous occasions from *E. coli*, first via the acquisition of the large ∼220 kb Mxi/Spa type 3 secretion system containing virulence plasmid and subsequently through further gene acquisition and loss. They are the cause of bacillary dysentery, a severe infection of the intestinal mucosa resulting in colonic epithelial destruction, severe diarrhea, intestinal hemorrhage and, on occasion, hemolytic uremic syndrome (45). There are four different species of *Shigella, S. flexneri, S. sonnei, S. boydii* and *S. dysenteriae*, which have varied geographical distributions and can be further divided into many serotypes, dependent on O-antigen characteristics and other biochemical differences (45). Responsible for ∼500K deaths annually, understanding how these bacteria have evolved and how they respond to host and environmental stressors may shed light on better ways to combat them.

Two-component signaling systems are crucial for allowing bacteria to respond to changing environmental conditions and integrate those signals into an appropriate phenotypic output, thereby allowing bacteria to colonize a particular niche. The PhoPQ-PmrD-PmrAB system of *S. enterica* is, by far, the best characterized system in the context of enteric pathogenesis (5, 46, 47). In spite of the conservation of these systems in enteric pathogens, there is substantial heterogeneity in how different microbes integrate signals into a specific output as well as what accessory genes fall under the specific control of PhoPQ and/or PmrAB.

PhoPQ responds to numerous environmental signals including lowered pH and the presence of cationic host-defense peptides - signals associated with the intracellular phagosomal/vacuolar environment, although the importance of each signal to virulence may not be equal (48). In contrast, PmrAB responds to high concentrations of ferric iron as well as high concentrations of bile salts, signals that are more commonly associated with the lumenal small intestinal environment, particularly in the duodenum where bile salts are found in high concentrations and ferric iron solubility is increased due to exposure to the low pH environment of the stomach and before it is reduced to ferrous iron by duodenal cytochrome B (41, 49). In the context of mucosal antimicrobial peptide protection, host defense peptide production is highest in the ileum, where production of α-defensins by Paneth cells and β-defensins by intestinal epithelial cells are highest but colonic sites are also protected by host-derived LL-37 (50, 51). This is consistent with PmrAB being necessary for high level resistance to HDPs at mucosal sites where enteric bacteria initially encounter HDPs.

In addition to playing a role in the regulation of host defense peptide/polymyxin B resistance, PhoPQ it is also a crucial regulator of virulence in many enteric pathogens, including *S. enterica* (52, 53), enterohemorrhagic *E. coli* (54) and avian pathogenic *E. coli* (55, 56) and in *S. flexneri* (5, 31). The specific wiring of PhoPQ/PmrAB regulation can vary significantly from microbe to microbe (57–59), with the PmrD protein playing an important role in connecting these two systems in some microbes. PmrD works by binding to phospho-PmrA and prevents dephosphorylation by the sensor kinase/phosphatase PmrB (30, 60). This leads to enhanced activation of PmrA-dependent promoters. As PmrD is itself transcriptionally controlled by the PhoPQ system, the net effect of PmrD function is to bring the output of the PmrAB signaling system under the control of the PhoPQ system (21). Orthologues of the *pmrD* gene are found in *Klebsiella, Salmonella, Escherichia* and *Shigella*, but are notably absent from other Enterobacterales, including *Yersinia spp*., *Citrobacter* spp., *Erwinia* spp., *Providencia* spp. and *Enterobacter* spp. In spite of this difference, genes that are PmrAB-dependent in *Escherichia* and *Salmonella*, like the *arnBCDTEFugd* operon, responsible for the addition of 4-aminoarabinose to the 1 or 4’ phosphate of lipid A, are often directly transcriptionally regulated by the PhoPQ and PmrAB systems of genera that lack PmrD, suggesting that there are multiple ways to integrate the PhoPQ and PmrAB signaling systems to produce the desired output (57, 61).

Canonical PmrAB regulated genes include many that are involved in alteration of the surface properties of the bacteria. These include alterations to lipid A 1 and 4’ phosphate residues via the addition of 4-aminoarabinose and/or phosphoethanolamine (24, 62), the addition of phosphoethanolamine to KDO and heptose phosphates in the core region (25, 63) and by increasing the length of the O-antigen itself (64–67). The net effect of these modifications is to greatly reduce the interaction of the bacterial envelope with antimicrobial HDPs and components of the complement pathway and increase resistance to these types of molecules (64–66).

Constitutive activation of PmrA leads to alterations in outer membrane structure that can sensitize bacteria to bile salts and bile salts themselves are direct activators of enteric PmrAB signaling, suggesting that bacteria must finely balance the amount of PmrA-mediated modifications on their surface (49, 68). In *Salmonella*, PmrAB signaling is necessary for virulence via the oral route but is dispensable for virulence via the peritoneal route, suggesting that PmrAB is not required for virulence per se, but rather for adaptation to the mucosal environment (62) or by modulating the immune response to the infecting microbe (69).

*Shigella* spp. are a polyphyletic group of enteropathogens that have evolved from non-pathogenic *E. coli* strains following the acquisition of a large virulence plasmid containing a T3SS and numerous T3SS effectors. Following this acquisition, further gene gain and gene loss can alter the behaviour of *Shigella* spp. to fine tune control of virulence. All *Shigella* spp. have lost the function of the *cadA* gene. CadA encodes a lysine decarboxylase enzyme and the product of this, cadaverine, is a potent inhibitor of *Shigella* enterotoxin (70, 71). Similarly, while the OmpT protein is an important virulence factor in *E. coli*, contributing to HDP resistance in EHEC and other pathovars, no *Shigella* spp. produce functional OmpT protein, as it’s expression interferes with the normal function of the IcsA actin recruitment protein on the *Shigella* spp. pole (72). *Shigella* spp. have also deleted either the *nadA* and/or *nadB* genes, responsible for the production of a small molecule inhibitor of virulence in *Shigella* (73). Genes that are selectively lost in some lineages, leading to enhanced pathogenicity have therefore been termed “antivirulence genes” or “black holes”, observable only by the effect their loss has on the surroundings (70).

Here, we show that within the enteric pathogen *S. flexneri*, there has been selection against functional PmrD. Although the genome of *S. flexneri* appears to have a full-length functional copy of the *pmrD* ORF, it contains a frameshift that alters the location of the ORF with respect to the SD sequence and renders this gene non-functional. This loss makes the bacterium susceptible to the model cationic antimicrobial peptide, polymyxin B in a manner consistent with this loss. In *S. flexneri*, PmrAB regulated polymyxin B remains intact in this lineage, suggesting that the bacterium can still respond to luminal signals leading to enhanced antimicrobial peptide resistance but that when the bacterium in in an intracellular location, PhoPQ mediated control of PmrAB-dependent phenotypes is maladaptive. This is consistent with the *pmrD* gene of *S. flexneri* inhibiting some aspect of virulence associated behaviour and further analysis of these strains will shed light on the mechanisms by which this might be so. Genomic analysis of other *Shigella* spp. shows that the *pmrD* gene has been either deleted or mutated in some lineages of both *S. dysenteriae* and *S. boydii*, suggesting that there might be active selection against functional PmrD-PmrAB signaling in *Shigella* spp. As this connector protein is well conserved in *E. coli* and *S. enterica*, this loss suggests that *S. flexneri* may no longer require the function of PmrD during it’s virulence program, or indeed, the presence of functional PmrD protein may abrogate virulence in some way. We speculate that the maintenance of the start-site array shown in Figure 2 might offer some advantage to the population, perhaps by allowing a subset of a growing population to maintain expression of functional *pmrD*, when the frame-shift is repaired or otherwise reverted.

In addition to the specific effects associated with loss of PmrD in *Shigella flexneri*, our results also suggest that subtle modifications in the location of the translational start site relative to the ribosome binding site can have profound effects on the behaviour of a given gene and that bacteria could modulate behaviour of some pathways via this mechanism. Automated gene annotation programs typically annotate based on the presence in the genome of an intact open reading frame with a specific level of homology to a previously characterized gene in another organism (74). Although these programs typically also consider the location of the RBS relative to the 5’ end of the gene, automated annotation programs have often had difficulty in correctly identifying the correct start codon for a given gene (75). Our results provide a specific example of what may be a more generalizable pattern in other regulatory or metabolic pathways. Further investigation of this possibility is warranted.

